# Interaction of surface topography and taper mismatch on head-stem modular junction contact mechanics during assembly in modern total hip replacement

**DOI:** 10.1101/2022.01.20.476985

**Authors:** Jonathan A. Gustafson, Steven P. Mell, Brett Levine, Robin Pourzal, Hannah J. Lundberg

**Author notes:** Address correspondence to Jonathan A. Gustafson, Ph.D. Rush University Medical Center, 1611 W Harrison St, Suite 204, Chicago, IL, 60612. USA.

## Abstract

Implant failure due to fretting corrosion at the head-stem modular junction is an increasing problem in modular total hip arthroplasty. The effect of varying microgroove topography on modular junction contact mechanics has not been well characterized. The aim of this work was to employ a novel, microgrooved finite element (FEA) model of the hip taper interface and assess the role of microgroove geometry and taper mismatch angle on the modular junction mechanics during assembly. A two-dimensional, axisymmetric FEA model was created using a modern 12/14 taper design of a CoCrMo femoral head taper and Ti6Al4V stem taper. Microgrooves were modelled at the contacting interface of the tapers and varied based on height and spacing measurements obtained from a repository of measured retrievals. Additionally, taper angular mismatch between the head and stem was varied to simulate proximal- and distal-locked engagement. Forty simulations were conducted to parametrically evaluate the effects of microgroove surface topography and angular mismatch on predicted contact area, contact pressure, and equivalent plastic strain. Multiple linear regression analysis was highly significant (p < 0.001; R^2^ > 0.74) for all outcome variables. The regression analysis identified microgroove geometry on the head taper to have the greatest influence on modular junction contact mechanics. Additionally, there was a significant second order relationship between both peak contact pressure (p < 0.001) and plastic strain (p < 0.001) with taper mismatch angle. These modeling techniques will be used to identify the implant parameters that maximize taper interference strength via large in-silico parametric studies.

## INTRODUCTION

Modularity in total hip replacement (THR) provides the surgeon with a high degree of flexibility in conducting the arthroplasty procedure. However, modularity also leaves additional contacting interfaces. One such interface that has been a site of fretting corrosion and subsequent THR failures occurs between the “female” head taper component and the “male” stem taper. The onset of fretting in this modular taper interface is mechanically driven by micromotion between the modular tapers in which wear debris is generated and can lead to an adverse local tissue reaction^1–6^. The failure rates of THRs due to fretting corrosion at the taper junction has been shown to be a consistent problem since the 90s, despite numerous changes by manufacturers to improve the tolerances of these two components^7–10^. The taper design has gone through considerable changes over the last three decades, with the 12/14 taper design being the most commonly used^11,12^. However, significant variability across manufacturers and even within each manufacturer’s 12/14 design may continue to convolute the problem of fretting corrosion^13^.

To ensure a stable locking interface between the femoral head and stem taper, microgrooves were introduced by manufacturers to increase the contact pressures within the modular junction and improve the interference fit upon assembly. However, it is not clear how the microgroove geometry—defined by the machining process—is chosen, as the variability in the microgroove geometry can vary across designs and even within the same implant design as has been noted in past retrieval work^13–15^. Any variability in the microgroove geometry could have substantial implications on the final interference fit of these components and, eventually, onset of fretting and corrosion. Our group has shown through retrieval analysis that the surface topography is related to damage scores and damage modes found on retrieved implants (both postmortem and revised implants)^14–18^. However, the role of taper surface topography in conjunction with additional confounding implant design parameters—such as relative taper mismatch angle—has seen limited investigation^19^. It is challenging for manufacturers to control the relative mismatch between the head and stem tapers; thus, microgrooves could provide some “mechanical resilience” by allowing mismatched tapers to elicit high contact pressures and maintain a secure fit. The relationship between the surface topography and mismatch on modular taper assembly requires additional investigation to optimize the taper fit under different implant designs.

Prior experimental work has been performed to understand taper design changes on assembly mechanics via large benchtop tests^20–23^. However, these benchtop tests are not optimal (both time and costs) for conducting large parametric studies to evaluate the potential effects of taper design changes on taper mechanics. Computational modeling is an excellent, complimentary tool to parametrically examine the effects of taper design on mechanics. Several groups have modeled the modular taper interface yet commonly ignore the role of surface topography for both the stem and head taper^19,24–27^. We have shown it is vital to include surface topography in models to replicate the plastic deformation found in retrievals^14,28,29^. Modeling the plastic deformation within the modular taper junction is critical to effectively approximate the mechanics occurring in vivo. Thus, the objective of this study was to employ a novel, microgrooved finite element (FEA) model of the hip taper interface and assess the role of microgroove geometry and taper mismatch angle on the modular junction mechanics during assembly.

## METHODS

### FINITE ELEMENT ANALYSIS

A two-dimensional, axisymmetric model of a CoCrMo femoral head taper and Ti6Al4V stem taper was created using median measurements taken from 100 retrieved implants^15^. All models represented a 12/14 taper design. Surface topography in the form of microgrooves on the stem and head taper were modeled using a sinusoidal function with amplitude and period corresponding to median retrieval measurements of height and spacing, respectively. The nodes located at the external surface of the head taper geometry were constrained and tied to a reference node for control of the geometry. The bottom of the stem taper was fixed, and symmetry was assumed in the radial direction for both head and stem geometries (**Figure 1**).

**Figure 1.**
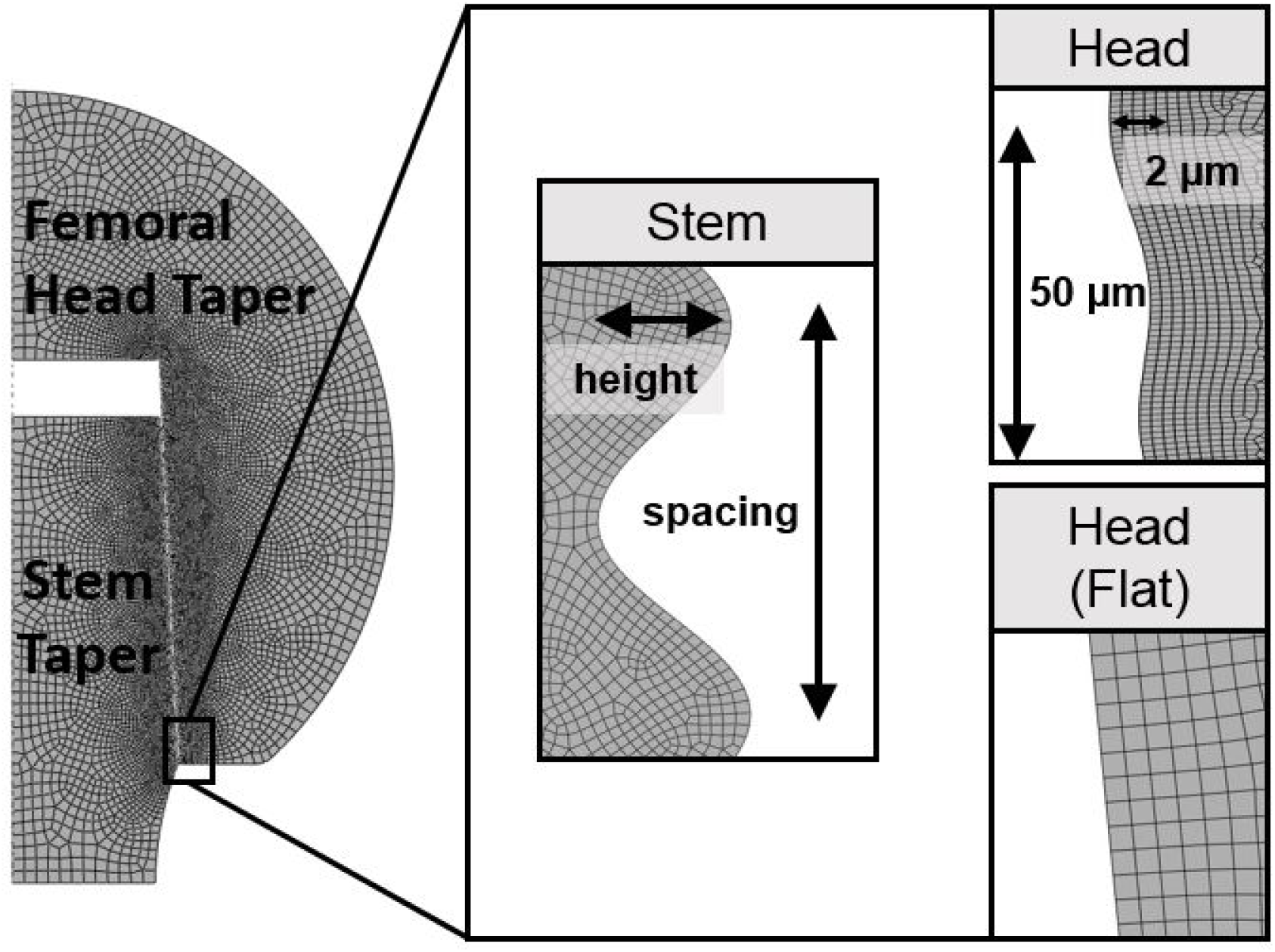
Finite element model of the head/stem modular taper (left) with representative modeling of microgroove surfaces (inset)

To simulate assembly during surgery, the head taper was moved onto the stem taper through control of the femoral head reference node at a constant velocity until a 4kN reaction load was achieved. The CoCr alloy was modelled as an elastic-linear plastic material with the following properties: Young’s modulus = 210 GPa, Poisson’s ratio = 0.30, yield strength = 827 MPa, ultimate tensile strength = 1000 MPa and elongation = 15%. The Ti6Al4V alloy was modelled as an elastic-linear plastic material with the following properties: Young’s modulus = 119 GPa, Poisson’s ratio = 0.30, yield strength = 795 MPa, ultimate tensile strength = 900 MPa and elongation = 15%. Contact between the head and stem taper interface was modelled using surface-to-surface discretization with a finite sliding formulation. A friction coefficient of 0.2^30^ was used in the penalty method for the contact model. Models were assembled and meshed in ABAQUS Standard (v 6.17) using four-node linear hexahedral, reduced integration elements (CAX4R). The head and stem models were meshed with approximately 321,864 and 307,191 elements, respectively.

### MODEL VERIFICATION

A mesh sensitivity analysis was conducted to evaluate the change in outcome variables with changing mesh density. Five additional simulations with varying mesh densities (10 micron, 5 micron, 2 micron, 1 micron, and 0.5 micron) were conducted at the perfect alignment position (see below) with the baseline stem taper topography (height = 11 μm, spacing = 200 μm) and the head taper surface with topography.

### EFFECT OF MICROGROOVE GEOMETRY & TAPER ANGLE

The objective of this study was to evaluate the role of stem/head taper microgroove geometry and taper mismatch angle on modular junction assembly mechanics. The range of taper microgroove geometry used in this parametric study can be found in Table 1 and were taken from median (baseline geometry: 11 μm height, 200 μm spacing) and standard deviation of 100 tapers measured in our retrieval database^15,18^. When modeling the stem taper, a “smooth” stem taper profile consisted of measurable microgroove geometry that was substantially lower in height and spacing compared to the “rough” profiles, which commonly had large heights and spacing. The head taper was modeled as either a) an “ideal” flat surface (i.e. no microgrooves), or b) a “baseline” surface, where the microgroove geometry profile consisted of median height (2 μm) and spacing (50 μm) from measured head tapers in our retrieval database^15,18^. To investigate the role of angular mismatch on changes in taper mechanics, the relative head and stem taper angles were varied to simulate distal, proximal, and perfectly aligned taper angles (Table 2). The range of values used in the current study assume normal manufacturing tolerances (±3’ mismatch) and extend out to values that have been found in our retrieval database (±12’). Forty simulations (5 mismatch angles x 2 head taper surface types x 4 stem taper microgroove geometries) were run. Outcome variables included contact area and contact pressure, equivalent plastic strain, and number of microgrooves undergoing plasticity.

**Table 1:** Range of taper topographies used for parametric analysis.

| Stem Taper | Smooth | Semi-rough | Baseline | Rough |
| --- | --- | --- | --- | --- |
| Height (μm) | 2 | 6 | 11 | 14 |
| Spacing (μm) | 30 | 150 | 200 | 200 |
| Head Taper | “Ideal” Flat |  | Baseline |  |
| Height (μm) | n/a |  | 2 |  |
| Spacing (μm) | n/a |  | 50 |  |

**Table 2:** Head and stem taper angles. Angular mismatch was calculated as the head taper angle – the stem taper angle. Degree-minutes-seconds notation is used to define the arcminute (‘) and arcsecond (“) for each taper mismatch by multiplying each taper angular mismatch by 60.

| Taper Mismatch | Distal Locked |  | Perfect fit | Proximal Locked |  |
| --- | --- | --- | --- | --- | --- |
|  | -12′ | -3′ | 0′ | 3′ | 12′ |
| **Head Taper Angle (°)** | 5.619° | 5.619° | 5.669° | 5.719° | 5.619° |
| **Stem Taper Angle (°)** | 5.819° | 5.669° | 5.669° | 5.669° | 5.419° |
| **Angular Mismatch (°)** | -0.2° | -0.05° | 0° | 0.05° | 0.2° |

The influence of mismatch angle and microgroove geometry on predicted outcomes from the model were evaluated using multiple linear regression in RStudio v1.4 (RStudio Team, Boston, MA). Stem and head taper topographies were modelled as categorical variables and levels are found in Table 2. Angular mismatch was considered a continuous variable and ranged from −12’ to 12’ (Table 2). All interaction terms were included in the analysis as well as the quadratic term for angular mismatch. ANOVA was used to test significance of each factor, and significance was set at the level of α = 0.05. Normality and homogeneity of the model were confirmed by Quantile-Quantile (Q-Q) plots and fitted value vs residual plots, respectively.

## RESULTS

### MODEL VERIFICATION

Results of the mesh sensitivity analysis showed less than a 3% change in total displacement, contact area, and contact pressure when going from 1 μm to 0.5 μm mesh density, verifying that a 1 μm mesh density was sufficient to capture the modular taper junction mechanics. The 1 μm mesh density also allowed for excellent approximation of the curvature of the stem taper microgrooves with varying topography parameters.

### EFFECT OF MICROGROOVE GEOMETRY & TAPER ANGLE

In general, changes in stem taper microgroove height led to substantial changes in both contact area and contact pressure predicted by the FEA model (Figure 2). Increasing stem taper microgroove height led to decreased contact area and subsequent increases in contact pressure and plastic strain at each microgroove in contact. Models with head taper microgroove topography exhibited substantially decreased contact area and conversely led to increased contact pressure and plastic deformation, whereas head taper models with a flat contact edge (i.e. “perfectly smooth”) exhibited the lowest contact pressures and no plastic deformation. A greater degree of angular mismatch resulted in contact initiating at the distal or proximal edge depending on the direction of mismatch and did not substantially change the contact mechanics in terms of contact area.

**Figure 2.**
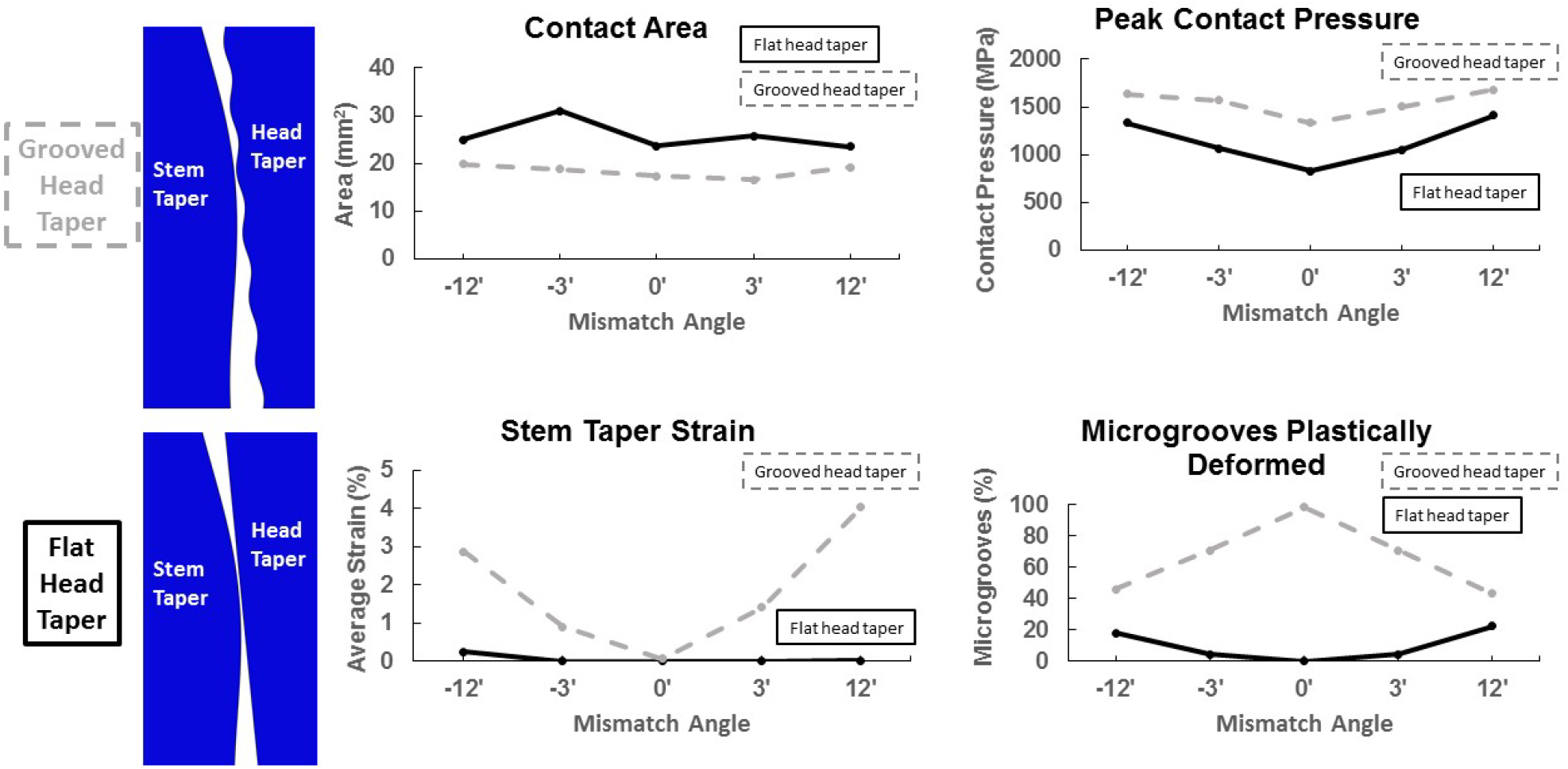
Example modular taper mechanics resulting from modeling an “ideal flat” head taper surface and modeling the baseline grooved head taper

A multiple linear regression model was used to determine the most important implant parameters contributing to model results of contact area, peak contact pressures, and peak plastic strain across the stem taper (Figure 3). Both peak contact pressure and peak plastic strain were calculated as the peak contact pressures or strains exhibited by each microgroove and averaged across all microgrooves. All regression models were highly significant (p < 0.001) and had R^2^ > 0.74. The regression analysis identified microgroove geometry on the head taper to have the greatest influence on modular junction contact mechanics, showing up as significant in all three regression models. When predicting the contact area between the stem and head taper, stem taper microgroove geometry (p = 0.0139) and the presence of head taper microgrooves (p < 0.001) were significant predictors. Mismatch angle was not a significant predictor of contact area within the modeled range. When predicting the peak contact pressure across the stem taper microgrooves, the presence of head taper microgrooves (p < 0.001) and stem taper topography (p < 0.001) were significant predictors. There was a significant second order relationship between peak contact pressure (p < 0.001) and mismatch angle. In addition, there was a significant interaction between the presence of head taper topography and stem taper topography (p =0.03). When predicting the peak microgroove plastic strain across the entire stem the presence of head taper microgrooves was a significant predictor (p < 0.001), as well as the second order term for angular mismatch (p < 0.001).

**Figure 3:**
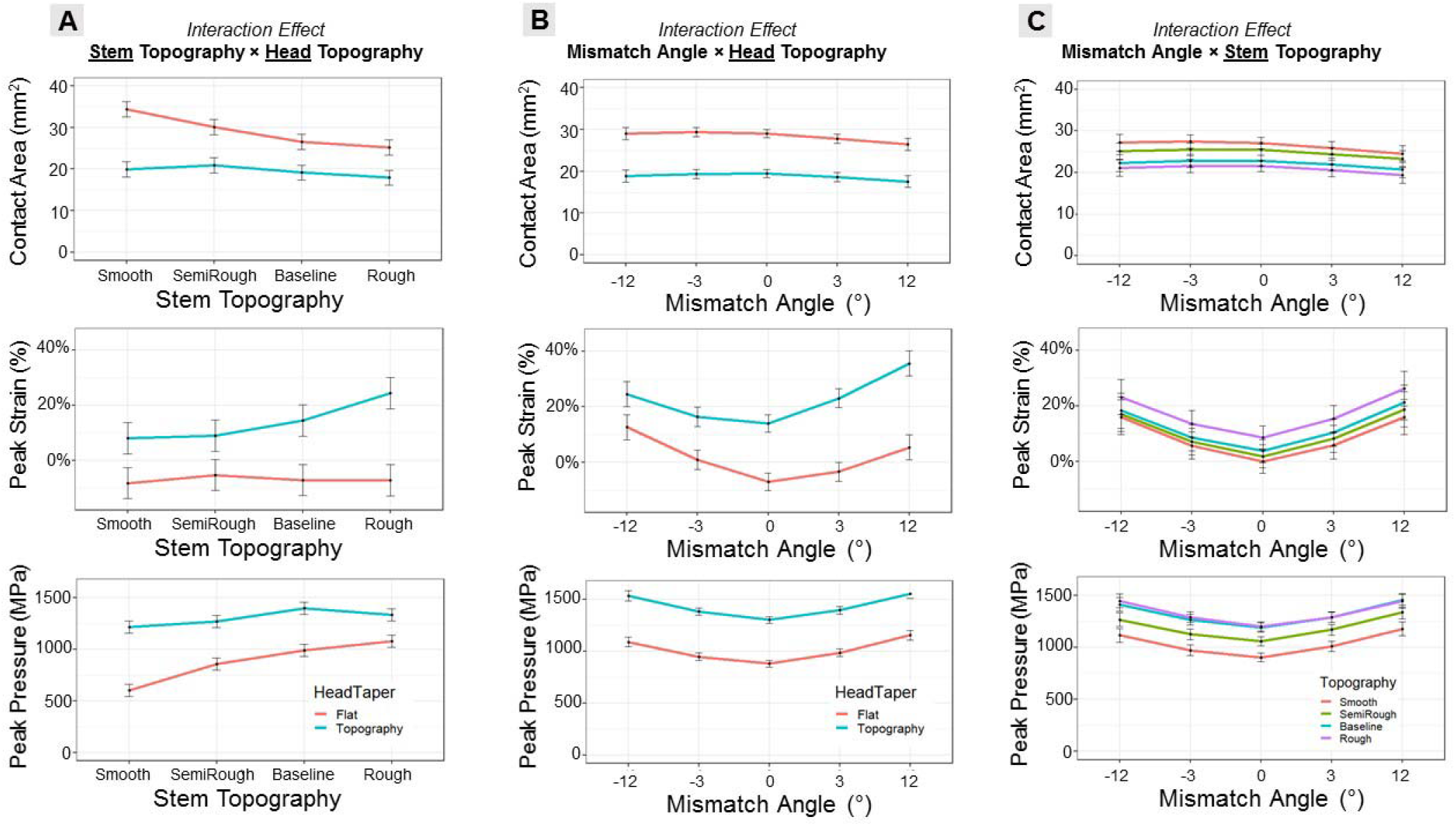
Results of the multiple linear regression model showing the interaction effects of a) stem taper topography by head taper topography, b) taper mismatch angle by head taper topography, and c) taper mismatch angle by stem taper topography on simulated contact area, peak strain, and peak contact pressures (individual rows).

## DISCUSSION

The contact mechanics of the modular taper junction are critical to its ability to stabilize the replacement. Alterations in the contact mechanics is of interest due to adverse local tissue reactions arising from the release of metal ions and particulate debris associated with fretting-corrosion at this interface. The objective of this study was to employ a novel, microgrooved FEA model of the hip taper interface to assess the role of microgroove surface topography and taper mismatch angle on the modular junction mechanics during assembly. We found that increased stem taper microgroove height led to reduced total contact area in the taper but greater contact pressures and plastic deformation. We also found substantial differences in modular contact mechanics when modeling the head taper with and without microgroove surface topography such that ignoring head taper surface topography (i.e. flat edge) led to no plastic deformation of the stem taper contact surface. Lastly, we found that taper angular mismatch exhibited a quadratic response on contact pressures and plastic strains across each taper, indicating increased contact pressures and plastic strain with increasing taper mismatch (both positive and negative mismatch).

The major finding of this study is that the presence of microgrooves on the stem and head taper significantly influence the contact mechanics predicted from FEA models. Prior modeling work has largely ignored the presence of microgrooves, and only recent work has started incorporating microgrooves on the stem ^19, 24–26^ but commonly model the head taper as “smooth” (i.e. flat). This study has demonstrated that the assumptions of a flat head taper substantially alter the contact mechanics, leading to reduced contact pressures and little to no plastic deformation across the stem taper. Plastic deformation of the stem tapers has consistently been found in retrieved implants ^17,31,32^ and is a clear mechanical response to the in vivo loading environment; thus, any computational models attempting to recreate the in vivo mechanisms must properly model the surface topography present at this interface to appropriately represent the real-world contact mechanisms. Even “smooth” taper designs consist of small asperities in their surface topography, which we represented via models consisting of short (2 μm) and narrow (30 – 50 μm) microgroove surface topography. The 2D-axisymmetric models employed in the current work have been validated by our group^28,29^. In the first study, contact mechanics were compared between FEA models and in vitro tests. In brief, CoCr head tapers were experimentally assembled onto Ti6Al4V stems at varying loads. Predicted microgrooves in contact (number) and those undergoing plastic deformation compared well to experimental changes across the loading conditions^29^. In our second study, we showed the change in microgroove height and width after assembly predicted by our FEA models were similar to changes in heights and spacing observed from retrieved implants with minimal implantation time^28^. While modeling the microgroove geometry is technically and computationally challenging, our work highlights the importance of accurately modeling this interface when researching questions relating to taper contact mechanics, extending to applications of fretting and wear.

Contact pressure was highly correlated to the input factors (R^2^ = 0.90). Based on the results of the regression model, a head with topography and stem taper topography at ‘baseline’ had the highest peak contact pressure. There was also a significant interaction between head taper topography and stem taper topography. In the case of a flat head taper, contact pressure increases as the stem taper goes from ‘smooth’ to ‘rough’. When topography is modelled on the head taper, contact pressure seems more resistant to changes in stem taper topography, with the resulting contact pressure plots being ‘flatter’ (Figure 3, column A). When predicting contact area, both head and stem taper topography were significant predictors, and the statistical model as a whole explained the variability in the data well (R^2^ = 0.76). The FEA models found that the greatest contact area was found on flat head tapers, as would be expected, with a smooth stem taper on a flat head having the highest contact areas. Based on the results of the linear regression, contact area on a head with topography appears to be somewhat resistant to changes in stem taper topography. Mismatch angle was not a significant predictor of contact area. When examining average peak plastic strain—representing the plastic deformation of the taper—topography on the head taper and the quadratic response for mismatch angle were significant predictors according to our linear regression. When investigating the results of our multiple linear regression analysis, the fitted values vs. residuals plot showed some degree of heteroscedasticity, potentially violating the assumptions of the linear regression. Despite the observed heteroscedasticity, the results of the ANOVA were a) consistent with previous results^14,15,29,33^ and b) make physical sense. A prior statistical model that was developed from 269 retrieved implants showed that increasing machined microgroove height led to decreased implant damage scores over implantation time^14^. The rationale for implementing microgrooves in the Morse Taper design was to allow for more effective collapse of the microgrooves under loading from impaction. As shown in our simulations and another benchtop study^34^, the peaks of the microgrooves undergo plastic strain under impact load and varies depending on the surface profile. For these reasons we are confident that we can conclude that the presence of topography on the head taper significantly affects plastic strain, and that the observed quadratic behavior of the effect of taper mismatch angle is valid.

One surprising finding was that taper angular mismatch did not substantially influence the contact mechanics of the modular junction, nor was it a major predictor of predicted mechanical outcomes from our stepwise regression model. Prior benchtop work^22,33^ and computational studies^24,25^ have assessed the effects of taper mismatch on general contact mechanics (i.e. not accounting for taper microgrooves) and commonly concluded that a more aligned taper would lead to improved interference strength. Our model predictions showed that the inclusion of microgroove surface topography on both the stem and head tapers were mechanically resilient to small taper mismatches (e.g. 3’ mismatch). Taper mismatch only seemed to influence the contact mechanics in the most extreme cases (i.e. 12’ mismatch), where there was a dramatic decrease in contact area and corresponding increase in both contact pressure and plastic strain. While we believe increased contact pressure and plastic strain is important to ensuring a stable interface, one confounding effect of increased taper mismatch is that the total number of microgrooves undergoing plastic deformation substantially decreases with increasing taper mismatch. This reduction in total microgroove deformation makes sense as the increased mismatch leads to localized stresses in one area of the taper (i.e. proximal or distal) and does not lead to even loading throughout the entire taper. Our study shows that the simulated microgroove geometry from tapers on the market are resilient to taper mismatch and that taper mismatch within the tolerances found in our retrievals is unlikely to be a culprit in the fretting-corrosion phenomenon, which has also been supported by various retrieval studies^6,35^.

Our model is not without limitations. We used median values from the retrieval database at our institute to identify range of microgroove geometry parameters for the model as opposed to modeling any combination of height and spacing parameters of microgroove geometry. This may limit the possible microgroove geometry designs to those from our institute and limits the generalizability to all implants used in THA. Our rationale was to investigate influence of existing microgroove geometry profiles found from retrieved implants and, thus, being manufactured and used in patients at our institute. Future work could perform a large design-of-experiments study to potentially identify an optimal combination of height/spacing parameters that is more “forgiving” of taper angular mismatch or other design tolerances. We also only modeled one material combination, titanium-alloy and cobalt chromium-alloy, and did not account for other potential manufacturing tolerances, such as out-of-roundness and surface waviness. However, the strength of our work was that our 2D model has been previously validated to accurately approximate the real permanent deformation of the taper microgrooves as measured from both retrievals^28^ and experimental tests^29^.

## CONCLUSION

We employed a novel, microgrooved FEA model of the hip taper interface to assess the role of microgroove surface topography and taper mismatch angle on modular junction mechanics. We have shown that the presence of microgrooves on the stem and head taper significantly influence the contact mechanics predicted by the FEA models, and that models without microgroove topography—particularly on the head taper—led to no plastic deformation of the stem taper contact surface. We also found that taper angular mismatch exhibited a quadratic response in predicted contact pressures and plastic strain, such that increasing contact pressure and strain was associated with increasing taper mismatch, regardless of mismatch direction (i.e. proximal or distally locked). The deformations of the microgrooves predicted by these models can be used to investigate the effects of numerous implant parameters (e.g. taper size, material, topography, etc.) via large in-silico parametric studies to identify the most important manufacturing parameters to improve taper interference strength and mitigate potential mechanically-based fretting corrosion.

